# ACE2-lentiviral transduction enables mouse SARS-CoV-2 infection and mapping of receptor interactions

**DOI:** 10.1101/2021.02.09.430547

**Authors:** Daniel J. Rawle, Thuy T. Le, Troy Dumenil, Kexin Yan, Bing Tang, Cameron Bishop, Andreas Suhrbier

## Abstract

SARS-CoV-2 uses the human ACE2 (hACE2) receptor for cell attachment and entry, with mouse ACE2 (mACE2) unable to support infection. Herein we describe an ACE2-lentivirus system and illustrate its utility for *in vitro* and *in vivo* SARS-CoV-2 infection models. Transduction of non-permissive cell lines with hACE2 imparted replication competence, and transduction with mACE2 containing N30D, N31K, F83Y and H353K substitutions, to match hACE2, rescued SARS-CoV-2 replication. Intranasal hACE2-lentivirus transduction of C57BL/6J mice permitted significant virus replication in lungs. RNA-Seq analyses illustrated that the model involves an acute inflammatory disease followed by resolution and tissue repair, with a transcriptomic profile similar to that seen in COVID-19 patients. Intranasal hACE2-lentivirus transduction of IFNAR^-/-^ and IL-28RA^-/-^ mice lungs was used to illustrate that loss of type I or III interferon responses have no significant effect on virus replication. However, their importance in driving inflammatory responses was illustrated by RNA-Seq analyses. We also demonstrate the utility of the hACE2-lentivirus transduction system for vaccine evaluation in C57BL/6J mice. The ACE2-lentivirus system thus has broad application in SARS-CoV-2 research, providing a tool for both mutagenesis studies and mouse model development.

**AUTHOR SUMMARY:** SARS-CoV-2 uses the human ACE2 (hACE2) receptor to infect cells, but cannot infect mice because the virus cannot bind mouse ACE2 (mACE2). We use an ACE2-lentivirus system *in vitro* to identify four key amino acids in mACE2 that explain why SARS-CoV-2 cannot infect mice. hACE2-lentivirus was used to express hACE2 in mouse lungs *in vivo*, with the inflammatory responses after SARS-CoV-2 infection similar to those seen in human COVID-19. Genetically modified mice were used to show that type I and III interferon signaling is required for the inflammatory responses. We also show that the hACE2-lentivirus mouse model can be used to test vaccines. Overall this paper demonstrates that our hACE2-lentivirus system has multiple applications in SARS-CoV-2 and COVID-19 research.

## INTRODUCTION

Severe acute respiratory syndrome coronavirus 2 (SARS-CoV-2) has spread rapidly into a global pandemic (1). SARS-CoV-2 infection can be asymptomatic, but is also the etiological agent of coronavirus disease 2019 (COVID-19), with acute respiratory distress syndrome (ARDS) representing a common severe disease manifestation (2). The SARS-CoV-2 pandemic has sparked unprecedented global research into understanding mechanisms of virus replication and disease pathogenesis, with the aim of generating new vaccines and treatments. Key to these efforts has been the development of animal models of SARS-CoV-2 infection and COVID-19 disease (3).

The receptor binding domain (RBD) of the spike glycoprotein of SARS-CoV-2 binds human Angiotensin-Converting Enzyme 2 (hACE2) as the primary receptor for cell attachment and entry (4). Mice do not support productive virus replication because the SARS-CoV-2 spike does not bind to mouse ACE2 (mACE2) (5). Expression of hACE2 in mice via a transgene allows SARS-CoV-2 infection and provides mouse models that recapitulate aspects of COVID-19. In such models hACE2 is expressed under control of various promoters, including K18 (6–12), mACE2 (13, 14), HFH4 (15), or chicken β-actin (16). These mouse models all differ in various aspects including level of virus replication, disease manifestations and tissue tropisms (3), but do not provide a simple mechanism whereby genetically modified (GM) or knock-out mice can be exploited for SARS-CoV-2/COVID-19 research. Two systems that do allow the latter, involve transient hACE2 expression in mouse lungs using adenovirus vectors (Ad5) or adeno-associated virus (AAV), which impart the capacity for productive SARS-CoV-2 replication in mouse lungs (10, 17, 18). A third model involves use of mouse adapted SARS-CoV-2 (19–21), although use of this mutated virus may complicate evaluation of interventions targeting human SARS-CoV-2.

Lentivirus-mediated gene expression in mice lung epithelium has been investigated as a treatment for cystic fibrosis and provides long term expression of the cystic fibrosis transmembrane conductance regulator (CFTR) (22–28). These studies demonstrated that VSV-G pseudotyped lentivirus can transduce mouse airway epithelial cells and their progenitors resulting in long-term gene expression. Herein we describe an ACE2-lentiviral system that conveys SARS-CoV-2 replication competence *in vitro* and allows productive infection of mouse lungs *in vivo.* We illustrate that GM mice can thereby be accessed for SARS-CoV-2/COVID-19 research, with experiments using IFNAR^-/-^ and IL-28RA^-/-^ mice showing the importance of type I and III IFN responses for SARS-CoV-2-induced inflammation. We also illustrate the use of hACE2-lentiviral transduction of lungs of wild-type C57BL/6J mice for evaluation of vaccines.

## RESULTS

### hACE2-lentiviruses for evaluating the role of RBD binding residues

hACE2 coding sequence (optimized for mouse codon usage) was cloned into the dual promoter lentiviral vector pCDH-EF1α-MCS-BGH-PGK-GFP-T2A-Puro (herein referred to as pCDH) (Figure 1A), which had been modified to contain the full length EF1α promoter. Previous studies have shown that the EF1α promoter effectively drives gene expression after intranasal VSV-G pseudotyped lentivirus transduction of mouse lungs (29). The GFP marker and puromycin resistance gene (cleaved from GFP by T2A) are expressed from a separate 3-phosphoglycerate kinase (PGK) promoter (Figure 1A). VSV-G pseudotyped pCDH-hACE2 lentivirus was produced by co-transfection of the lentiviral vector with VSV-G and Gag-Pol-Rev plasmids in HEK293T cells, and was used to transduce HEK293T (human embryonic kidney) cells followed by puromycin selection. HEK293T cells do not express hACE2 (30) and therefore do not support SARS-CoV-2 replication, whereas HEK293T cells transduced with hACE2 supported significant virus replication (Figure 1B). A T92Q mutation in hACE2 was introduced to determine if removing the N90 glycosylation motif increased SARS-CoV-2 replication, as computer-based modelling predicted enhanced affinity for spike RBD with removal of this glycan (31, 32). We also introduced K31N/K353H and T27Y/L79Y/N330Y mutations into hACE2 as these changes were predicted by computer modeling to reduce and enhance affinity for spike RBD (31), respectively. HEK293T cells expressing these aforementioned hACE2 mutations did not significantly affect SARS-CoV-2 replication (Figure 1B), illustrating that *in silico* affinity predictions do not always translate to differences in virus replication.

**Figure 1.**
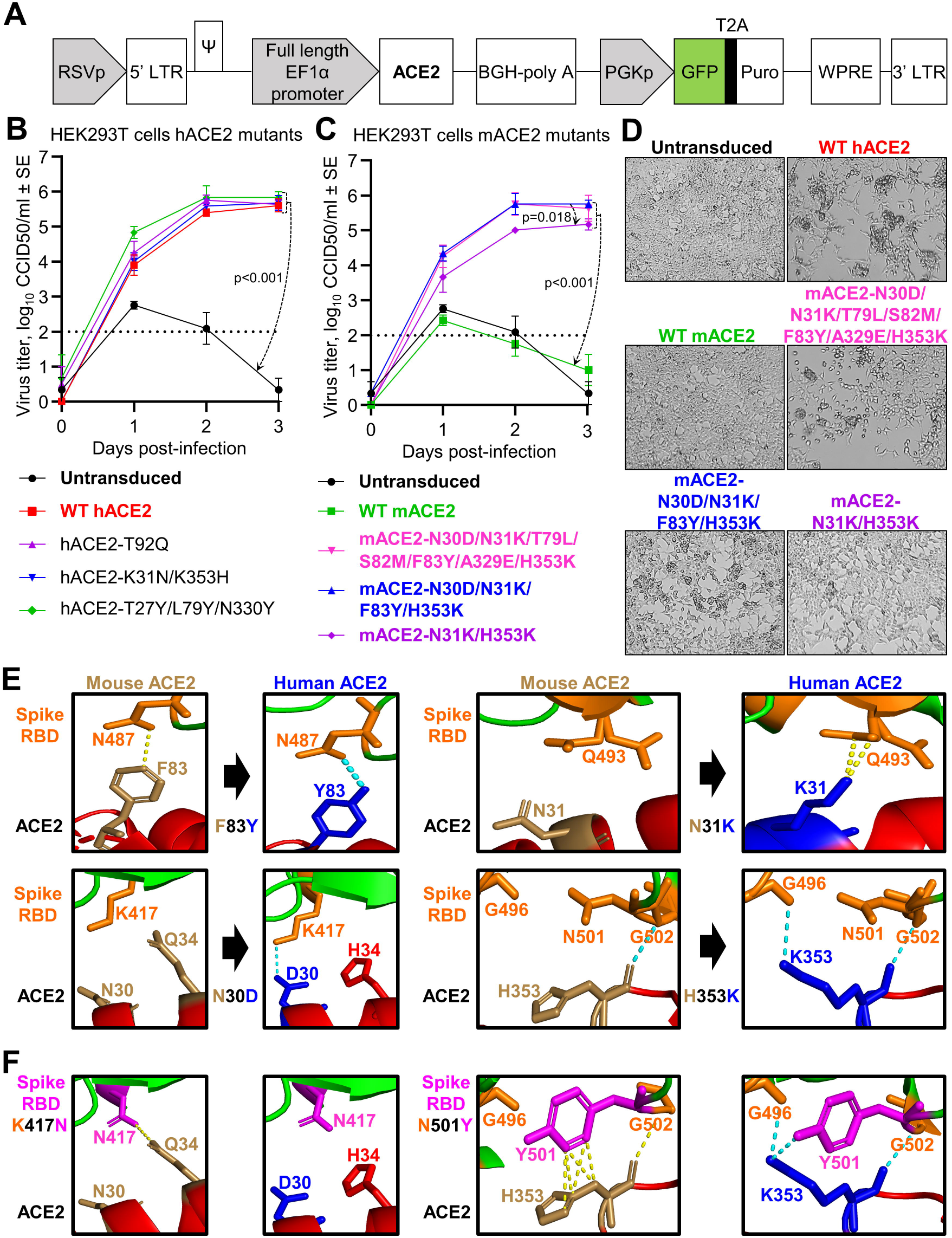
Mutational analyses of human and mouse ACE2 using ACE2-lentiviruses *in vitro*. **A)** Schematic of pCDH-EF1α-ACE2-BGH-PGK-GFP-T2A-Puro lentiviral vector. **B-C)** Growth kinetics of SARS-CoV-2 over a three day time course in HEK293T cells transduced with WT or mutant hACE2 **(B)** or mACE2 **(C)** infected at MOI=0.1. Data is the mean of 1 (hACE2-T27Y/L79Y/N330Y and mACE2-N31K/H353K) or 2 (all others) independent experiments with 2-3 replicates in each and error bars represent SEM. Statistics was determined using Repeated Measures Two-Way ANOVA comparing untransduced (for B) or WT mACE2 (for C) with all others, and comparing mACE2-N30D/N31K/F83Y/H353K with mACE2-N31K/H353K. **D)** Inverted light microscopy images of HEK293T cells transduced with the indicated ACE2 lentivirus and infected with SARS-CoV-2 at MOI=0.1. Images were taken at day 3 post infection and were representative of triplicate wells. **E)** Crystal structure of the spike RBD:hACE2 complex (PBD: 6M0J) (95) viewed in PyMOL and zoomed in on key interactions between ACE2 residues identified in ‘C-D’ (hACE2 = blue, mACE2 = brown) and spike RBD residues (orange). Yellow dotted lines represent any contacts between chains within 3.5Å, and blue dotted lines represent polar contacts. Direction of black arrows indicate predicted enhanced interactions. **F)** K417N and N501Y mutations (magenta) were introduced in the spike RBD to mimic the Republic of South Africa (B.1.351 and 501Y.V2) and United Kingdom (B.1.1.7 and 501Y.V1) SARS-CoV-2 variants.

### Characterizing residues responsible for the inability of mACE2 to support SARS-CoV-2 replication

The ACE2-lentivirus system was used to introduce amino acid substitutions into mACE2 to identify the amino acids that are responsible for restricting SARS-CoV-2 replication in mACE2-expressing cells. Analysis of crystal structures and species susceptibilities has suggested residues that may be responsible for the differences in binding of the RBD to hACE2 and mACE2 (33–37). Based on these *in silico* studies, we identified seven residues potentially responsible for the human-mouse differences. mACE2-lentivirus vectors were then generated that contained all seven substitutions (N30D/N31K/T79L/S82M/F83Y/A329E/H353K), with two further vectors constructed to identify the minimum requirements for SARS-CoV-2 replication. The latter contained a subset of the seven changes; four amino acid changes (N30D/N31K/F83Y/H353K) and two amino acid changes (N31K/H353K).

As expected HEK293T cells expressing mACE2 did not support productive SARS-CoV-2 replication, and CPE was not observed. In contrast, all three mACE2 mutants significantly rescued virus replication and infection resulted in CPE (Figure 1C-D). However, virus replication was significantly lower for cells expressing mACE2 with only two substitution (N31K/H353K) (Figure 1C) and infection produced less CPE (Figure 1D). Thus two amino acid changes (N31K and H353K) in mACE2 were sufficient to support SARS-CoV-2 replication; however, at least four changes (N30D/N31K/F83Y/H353K) were required for virus replication and CPE to be comparable to cells expressing hACE2. These studies highlight the utility of the ACE2-lentivirus system for studying the role of ACE2 residues in productive SARS-CoV-2 infections.

### Modeling the RBD:ACE2 key interactions for mouse, human and virus variants

Of all the potential interactions between the RBD and ACE2 previously identify by X ray crystallography and of all the amino acid differences between human and mouse ACE2 (33, 37), our data identified a key role for 4 ACE2 mutations for conferring SARS-CoV-2 infectivity on mACE2 expressing cells. Modeling of the interactions between these 4 mutations and the RBD predict improved interactions between the virus and the receptor (Figure 1E), providing an explanation for the gain of function for the mutated mACE2. The F83Y substitution replaced a non-polar interaction between mACE2-F83 and RBD-N487, with a polar interaction between hACE2-Y83 and RBD-N487. The N31K substitution created new interactions between ACE2-K31 and RBD-Q493. The N30D substitution created a salt bridge interaction between ACE2-D30 and RBD-K417. Lastly, the H353K substitution created a new hydrogen bond with RBD-G496.

Recently, SARS-CoV-2 variants have emerged with mutations in the RBD, with two of the key mutations involving interactions with hACE2 residues 30 and 353 (38). Republic of South Africa (RSA) variants (B.1.351 and 501Y.V2) have the RBD mutations K417N and N501Y, and the United Kingdom (UK) variants (B.1.1.7 and 501Y.V1) have the N501Y mutation (38). Modelling predicts that the K417N mutation would result in the gain of an interaction between RBD N417 and mACE2-Q34, whereas the interaction between K417 and hACE2-D30 would be lost (Figure 1F). The selection for RBD-K417N in humans may therefore be more related to antibody escape (39, 40), rather than improving interactions with hACE2. The N501Y mutation in the RBD is predicted to generate new interactions between Y501 with mACE2-H353, and new polar interactions between Y501 and hACE2-K353 (Figure 1F). Our modeling supports the observation that viruses with K417N and N501Y mutations have increased affinity for mACE2, with both appearing in mouse-adapted SARS-CoV-2 (41).

### hACE2-lentivirus transduction of mouse cell lines imparts SARS-CoV-2 replication competence

To determine whether hACE2-lentivirus transduction could make mouse cell lines SARS-CoV-2 replication competent and thus available for SARS-CoV-2 research, 3T3 (mouse embryonic fibroblasts) and AE17 (mouse lung mesothelioma cell line) cells were transduced with hACE2. Significant virus replication was seen in transduced cells (Supplementary Figure 1), although somewhat lower than that seen in transduced HEK293T cells. Overt CPE was not seen (Supplementary Figure 1). This illustrates that the hACE2-lentivirus system can be used for mouse cell lines, but that the efficiency of viral replication may be cell line or cell type dependent.

### SARS-CoV-2 replication in hACE2-lentivirus transduced C57BL/6J, IFNAR^-/-^ and IL-28RA^-/-^ mice lung

To illustrate the utility of the hACE2-lentivirus system for use in GM mice, we investigated the role of type I and type III IFN receptor signaling in SARS-CoV-2 infections. C57BL/6J, IFNAR^-/-^ and IL-28RA^-/-^ were inoculated intranasally (i.n.) with 1.2-2.2×10^4^ transduction units of hACE2-lentivirus, equivalent to approximately 190-350 ng of p24 per mice (Figure 2A). An hour prior to administration of lentivirus, mice were treated i.n. with 1% lysophosphatidylcholine (LPC) to enhance transduction efficiency (25, 42). One week later mice were challenged with SARS-CoV-2 (10^5^ CCID_50_ i.n. per mice), and lungs were collected at 2, 4 and 6 days post infection for C57BL/6J and IFNAR^-/-^ mice and day 2 for IL-28RA^-/-^ mice (Figure 2A). Mice did not display overt clinical symptoms or weight loss (Figure 2B), consistent with some studies using Ad5-hACE2 (10), but not others (43). RT-qPCR analyses indicated similar levels of hACE2 mRNA after transduction of mouse lungs in each of the 3 mouse strains, with expression maintained over the 6 day course of the experiment at levels significantly higher than untransduced mice (Figure 2C).

**Figure 2.**
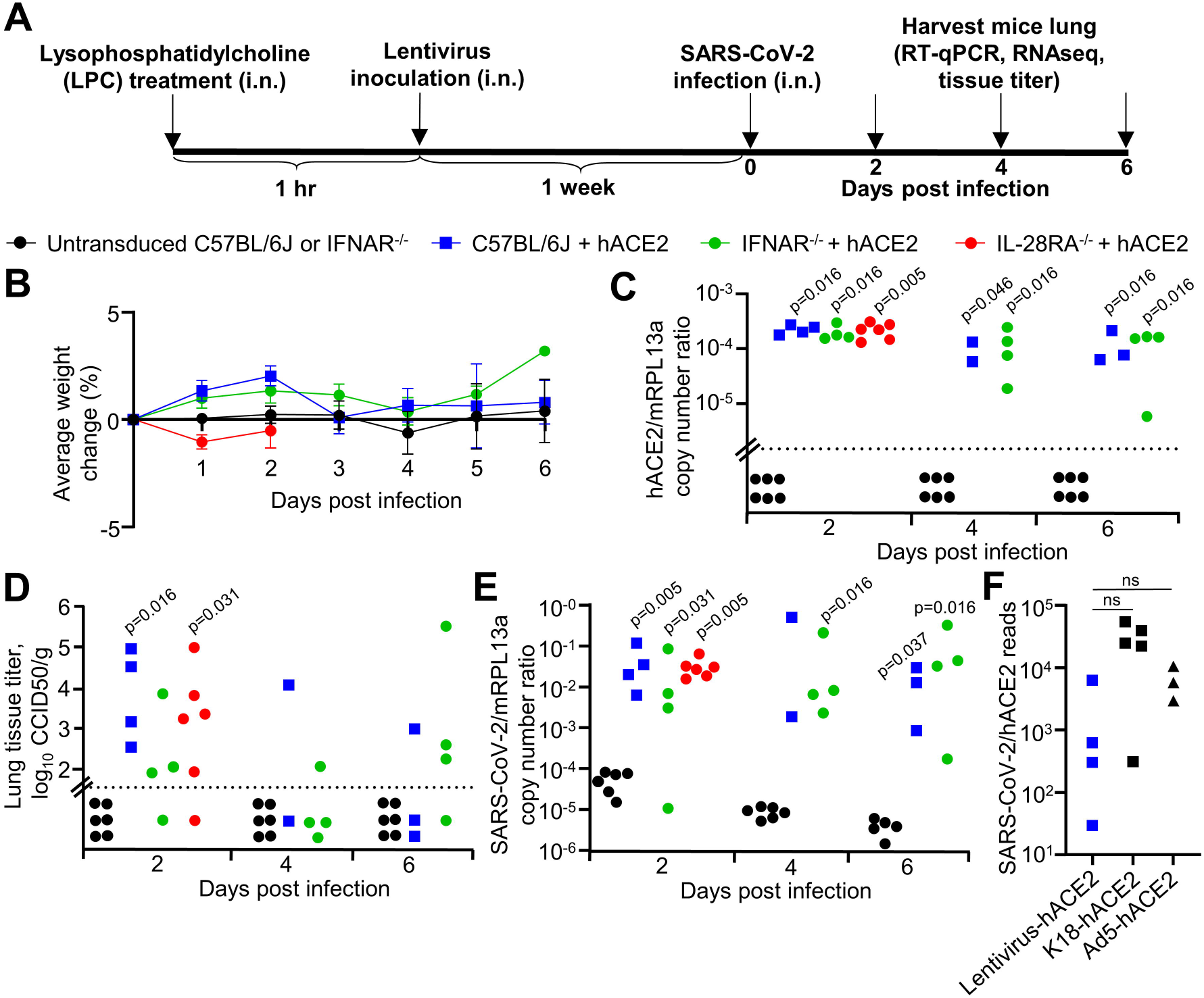
SARS-CoV-2 replication in hACE2-lentivirus transduced C57BL/6J, IFNAR^-/-^ and IL-28RA^-/-^ mice lung. **A)** Timeline of lentivirus transduction of mice lung and SARS-CoV-2 infection. **B)** Percent weight loss for mice in C-E. Data represents the mean percent weight loss from day 0 and error bars represent SEM. **C)** RT-qPCR of mice lung RNA using primers for hACE2 (introduced by lentivirus transduction) normalized to mRPL13a levels. Data is for individual mice and is expressed as RNA copy number calculated against a standard curve for each gene. Horizontal line indicates cut-off for reliable detection, with all untransduced mice falling below this line. Statistics are by Kolmogorov Smirnov test compared to untransduced samples. **D)** Titer of SARS-CoV-2 in mice lung determined using CCID_50_ assay of lung homogenate. Horizontal line indicates the limit of detection of 1.57 log_10_ CCID_50_/g. Statistics are by Kolmogorov Smirnov test compared to untransduced mice. IFNAR^-/-^ versus untransduced mice at day 2 post infection reaches significance by Kruskal-Wallis test (p=0.018). **E)** RT-qPCR of mice lung RNA using primers for SARS-CoV-2 E gene normalized to mRPL13a levels. Data is individual mice and is expressed as RNA copy number calculated using a standard curve. Statistics are by Kolmogorov Smirnov test compared to untransduced mice. C57BL/6J hACE2-transduced versus untransduced mice at day 4 post infection reaches significance by Kruskal-Wallis test (p=0.046). See Supplementary Figure 2 for SARS-CoV-2 RNA copies normalised to hACE2 copies. **F)** SARS-CoV-2 read counts (also see Supplementary Figure 3) normalized to hACE2 read counts from RNA-seq data for lentivirus-hACE2 transduced mice at day 2, K18-hACE2 transgenic mice at day 4, and Ad5-hACE2 transduced mice at day 2 (43). Not significant by Kolmogorov Smirnov or Kruskal-Wallis tests.

Lungs from infected C57BL/6J, IFNAR^-/-^ and IL-28RA^-/-^ were analyzed for infectious virus tissue titers. Untransduced lungs showed no detectable virus titers (Figure 2D), whereas hACE2-lentivirus transduced lungs showed significant viral titers ranging from 10^2^ to 10^5^ CCID_50_/g for all strains of mice on day 2, with titers dropping thereafter (Figure 2D). Viral RNA levels were measured using RT-qPCR, with untransduced lungs showing low and progressively decaying levels post-inoculation (Figure 2E). For all hACE2-lentivirus transduced lungs, significantly elevated and persistent viral RNA levels were seen for all strains and at all time points by RT-qPCR (Figure 2E). Reducing viral titers and lingering viral RNA have been reported previously for K18-hACE2 mice (9) and has also been suggested in human infections (44). RNA-Seq also illustrated that virus replication in hACE2-lentivirus transduced lungs, K18-hACE2 transgenic mice and Ad5-hACE2 transduced lungs (43) were not significantly different when normalized to hACE2 mRNA expression (Figure 2F), suggesting comparable hACE2 translation efficiencies and SARS-CoV-2 replication efficiencies in these different expression systems.

Importantly, no significant differences in viral loads emerged between C57BL/6J, IFNAR^-/-^ and IL-28RA^-/-^ mice (Figure 2D, E). This remained true even when viral RNA levels were normalized to hACE2 mRNA levels (Supplementary Figure 2). Similar results were obtained by RNA-Seq, with viral read counts not significantly different for the 3 mouse strains (Supplementary Figure 3A-D). Thus using three different techniques, no significant effects on viral replication could be seen in mice with type I or type III IFN receptor deficiencies.

### SARS-CoV-2 replication in hACE2-lentivirus transduced mice lung induces inflammatory signatures by day 2 post-infection

To determine the innate responses to SARS-CoV-2 replication in hACE2-transduced mouse lungs, gene expression in lungs of infected hACE2-transduced mice was compared with infected untransduced mice on day 2 post-infection using RNA-Seq. Differentially expressed genes (DEGs) between these groups were identified using a false discovery rate (FDR, or q value) threshold of <0.05 (Figure 3A; Supplementary Table 1A-C). Of the 110 DEGs, 95 were upregulated, with ≈ type I IFN-stimulated genes (ISGs) as classified by Interferome (using conservative settings that only included genes validated *in vivo* for mice) (Figure 3B, full list in Supplementary Table 1D). Type III ISGs are known to be poorly annotated in this database, and so this analysis likely under-estimates the number of such ISGs.

**Figure 3.**
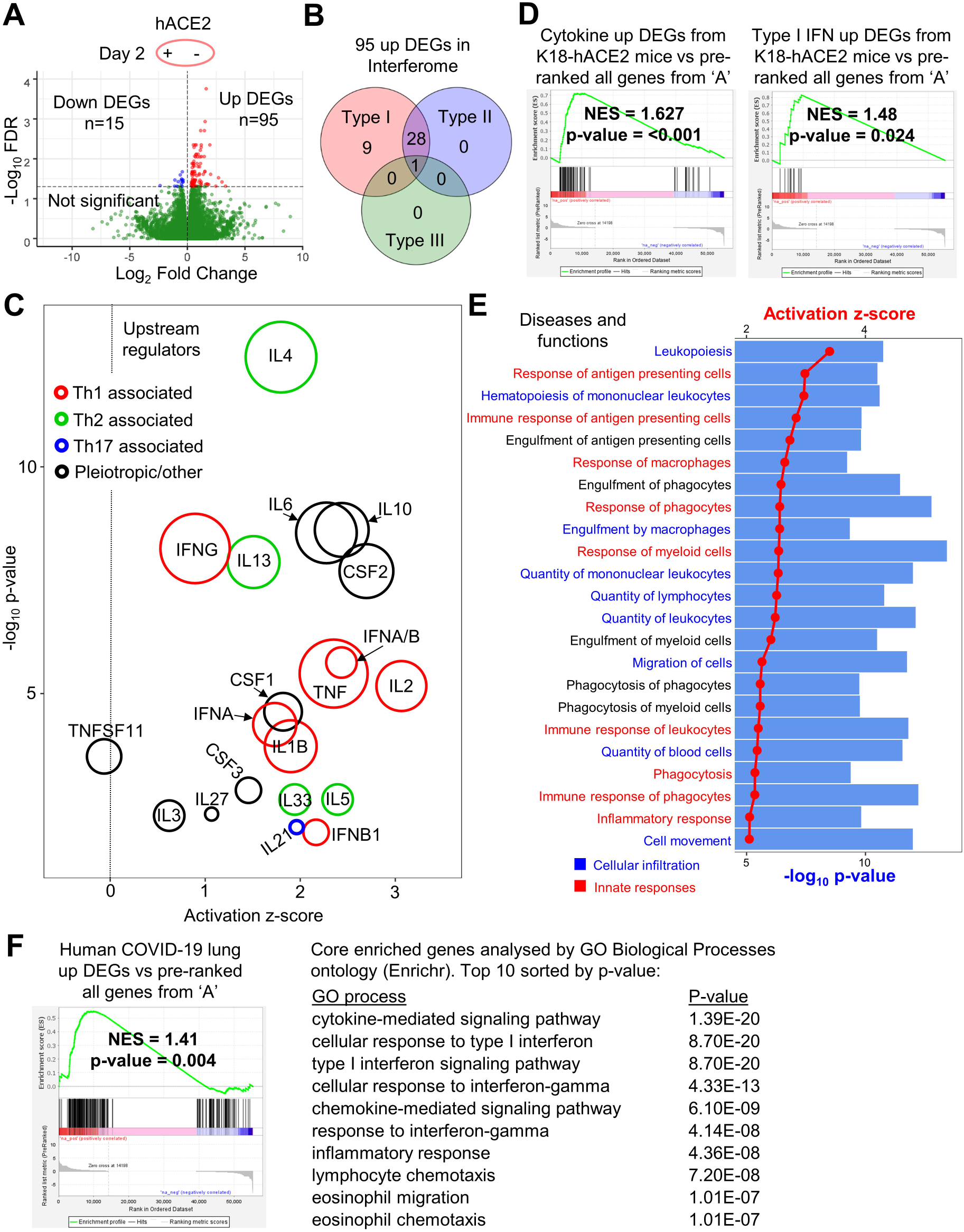
SARS-CoV-2 replication in hACE2-lentivirus transduced mice lung induces inflammatory signatures by day 2 post-infection. **A)** Volcano plot of gene expression from RNA-seq analyses of lung at day 2 comparing mice with and without hACE2-lentivirus transduction. Genes were considered DEGs if they had a FDR value < 0.05 (above horizontal dotted line) (see Supplementary Table 1B-C for full gene and DEG lists). **B)** Interferome analysis of 95 up DEGs from ‘A’. 38 of 95 up DEGs (40%) were ISGs. Red = type I IFN, blue = type II IFN, green = type III IFN (see Supplementary Table 1D for full Interferome ISG list). **C)** Cytokine signatures identified by IPA USR analysis (see Supplementary Table 1E-F for full and cytokine only USR lists) of 110 DEGs identified in ‘A’. Circle area reflects the number of DEGs associated with each USR annotation. USRs were grouped into three categories; red = Th1 associated, green = Th2 associated, blue = Th17 associated, and black = pleiotropic/other. The vertical dotted line indicates activation z-score of 0. **D)** GSEAs for cytokine-related DEGs or type I IFN-related DEGs from Winkler et al. supplemental data (9) against the pre-ranked all gene list (Supplementary Table 1B). Normalised enrichment score (NES) and nominal p-value are shown. **E)** IPA diseases and functions analysis (see Supplementary Table 1G-H for full lists) of 110 DEGs identified in ‘A’. The 23 annotations with the most significant p-value and with an activation z-score of >2 were plotted with dots/lines indicating activation z-score and bars indicating –log_10_ p-value. Annotations were grouped into two categories; cellular infiltration (blue) and innate responses (red). **F)** GSEA for DEGs from human COVID-19 lung versus healthy control from Blanco-Melo et al. (51) against the pre-ranked (by fold change) all gene list (Supplementary Table 1B). Normalised enrichment score (NES) and nominal p-value are shown. Core enriched genes (see Supplementary Table 1L for full core enriched gene list) determined by GSEA were entered into Enrichr and the top 10 GO Biological Processes annotations sorted by p-value are shown (see Supplementary Table 1M for full GO processes list).

Ingenuity Pathway Analysis (IPA) of the 110 DEGs produced a series of pro-inflammatory Upstream Regulators (USRs) (Supplementary Table 1E-F) and included Th1, Th2 and Th17-associated cytokine USRs, as well as type I IFN USRs (Figure 3C). SARS-CoV-2-specific T cells in humans were reported to be predominantly Th1-associated, but Th2-associated cytokines were also identified (45). A cytokine and type I IFN DEG list has been published for a RNA-Seq analysis of SARS-CoV-2 infected K18-hACE2 transgenic mice (9). These DEG lists were used to interrogate the full pre-ranked (fold change) gene list (Figure 3A, Supplementary Table 1B) by Gene Set Enrichment Analysis (GSEA). Significant enrichments were observed (Figure 3D), indicating similar cytokine and type I interferon (IFN) responses after SARS-CoV-2 infection of K18-hACE2 transgenic and hACE2-lentivirus transduced mice.

IPA *Diseases and Functions* analysis of the 110 DEGs revealed a series of annotations dominated by cellular infiltration and innate responses signatures (Figure 3E, Supplementary Table 1G-H). Several of the annotations were associated with monocytes and macrophages (Figure 3E), consistent with a previous report showing that inflammatory monocyte-macrophages were the major source of inflammatory cytokines in mouse adapted SARS-CoV infected mice (46). The cellular infiltrates in COVID-19 patient lungs is also dominated by monocyte-macrophages with a suggested role in severe disease (47). Innate immune signaling was also the dominant signature in other analyses including GO Biological Processes and Cytoscape (Supplementary Table 1I-J).

GSEA analysis using the >31,000 gene signatures available from the molecular signatures database (MSigDB), indicated significant negative enrichment (negative NES scores) of gene sets associated with translation and mitochondrial electron transport chain function (Supplementary Table 1K). Translation inhibition by SARS-CoV-2 Nsp1 via the blocking of mRNA access to ribosomes has been reported previously (48, 49). SARS-CoV-2 down-regulation of genes associated with cellular respiration and mitochondrial function has also been shown in human lung (50).

DEGs from human COVID-19 lung (51) were also enriched in our hACE2-lentivirus transduced mouse lungs, and analysis of the core enriched genes indicated the similarity was due to cytokine, IFN, chemokine and inflammatory signaling genes (Figure 3F, Supplementary Table 1L-M). Another human COVID-19 lung dataset was also interrogated (52) and the results again illustrated enrichment of immune-related DEGs in the hACE2-lentivirus transduced mice (Supplementary Table 1N).

Overall these analyses illustrate that SARS-CoV-2 infection in hACE2-lentivirus transduced C57BL/6J mice lungs significantly recapitulate responses seen in other murine models and in COVID-19 patients.

### Day 6 post-infection is characterized by inflammation resolution and tissue repair signatures

To determine how response profiles progress in the hACE2-lentivirus model, RNA-Seq was used to compare lungs on day 2 with lungs on day 6 post-infection, with day 6 broadly representing the time of severe lung disease in the K18-hACE2 model (9). As noted above (Figure 2D), the virus titers had dropped in hACE2-lentivirus transduced mice lung over this time period by ≈ 2.8 logs, and severe disease (typically measured by weight loss) was not evident in this hACE2-lentivirus model (Figure 2B). In both hACE2-lentivirus transduced and untransduced C57BL/6J mice there was a clear evolution of responses from day 2 to day 6 post-infection; however, DEGs obtained from the former were largely distinct from DEGs obtained from the latter (Figure 4A). The RNA-Seq data thus illustrated that the DEGs associated with virus infection were distinct from those associated with virus inoculation (Figure 4A). RNA-Seq of infected lungs provided 551 DEGs, 401 up-regulated and 150 down-regulated (Figure 4B, Supplementary Table 2A-C).

**Figure 4.**
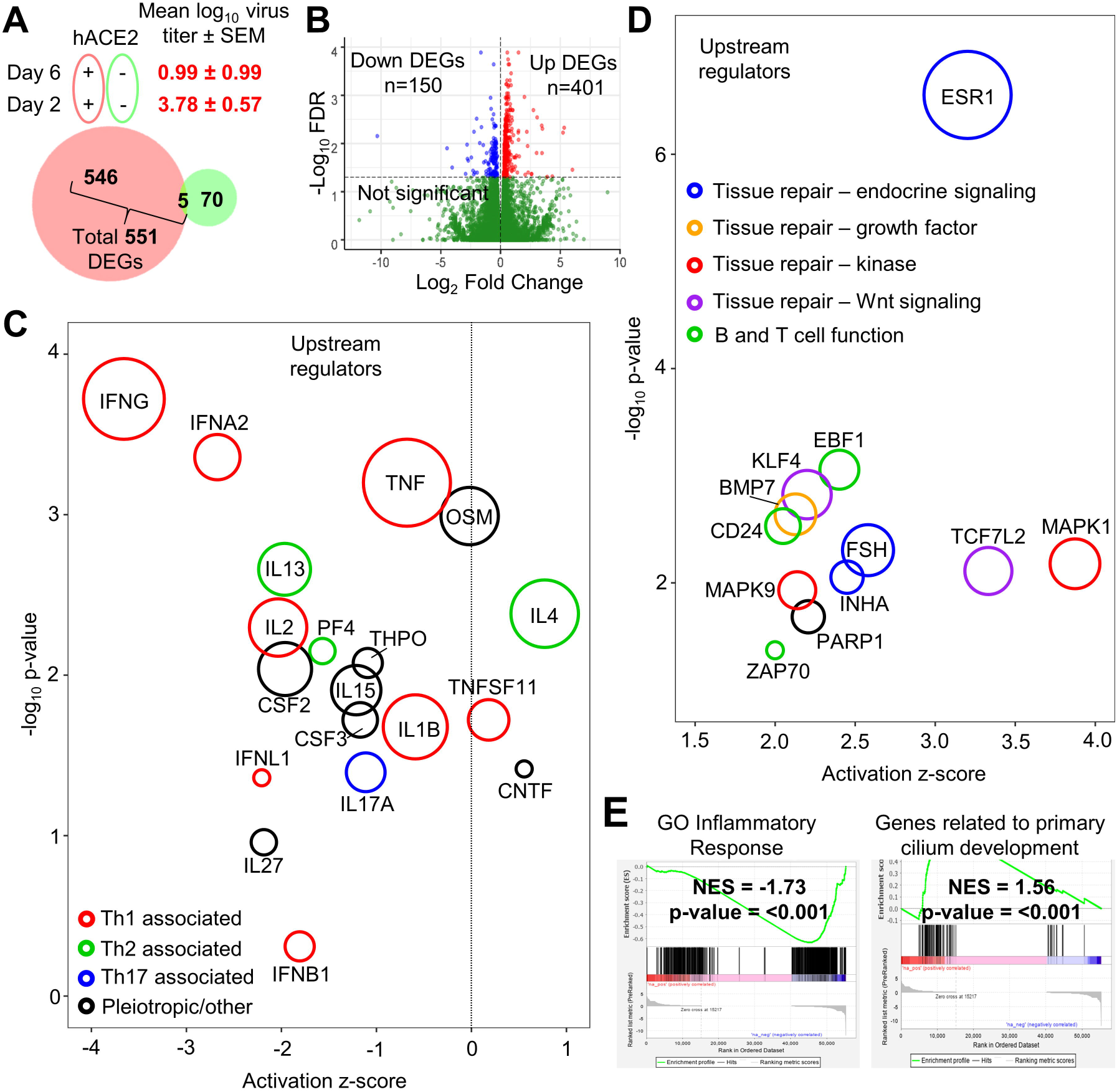
Day 6 post-infection is characterized by inflammation resolution and tissue repair signatures. **A)** Venn diagram for DEGs comparing day 6 versus 2 post infection with (red) and without (green) hACE2-lentivirus transduction. Number of DEGs that are FDR<0.05 are shown. Mean virus titer (log_10_ CCID_50_/g) in lung tissue ± SEM (represents data shown in Figure 2D) is shown for hACE2 transduced mice at day 2 or 6 post infection. **B)** Volcano plot of gene expression from RNA-seq analyses of hACE2-lentivirus transduced lung RNA comparing day 6 with day 2 (see Supplementary Table 2B-C for gene and DEG lists). Genes were considered DEGs if they had a FDR value < 0.05 (above horizontal dotted line). **C)** Cytokine signatures identified by IPA USR analysis (see Supplementary Table 2D-E for full and cytokine only lists) of 551 DEGs identified in ‘B’ (Supplementary Table 2C). Circle area reflects the number of DEGs associated with each USR annotation. USRs were grouped into four categories; red = Th1 associated, green = Th2 associated, blue = Th17 associated, and black = pleiotropic/other. The vertical dotted line indicates activation z-score of 0. **D)** Upregulated USR signatures identified using IPA analysis (see Supplementary Table 2F) of 551 DEGs identified in ‘B’. Circle area reflects the number of DEGs associated with each USR annotation. USRs (excluding chemicals) with an activation z-score of >2 are shown and were grouped into five categories; green = B cell function, blue = tissue repair – endocrine signaling, orange = tissue repair – growth factor, red = tissue repair – kinase, purple = tissue repair – Wnt signaling. **E)** Selected plots from GSEA of entire MSigDB database using pre-ranked (by fold change) ‘all’ gene list (Supplementary Table 2B). NES score and p-value is shown for the “GO_inflammatory_response” and “Wp_Genes_Related_To_Primary_Cilium_Development_Based_On_Crispr” annotations.

IPA USR analyses of the 551 DEGs showed a clear reduction in inflammatory cytokine annotations, although the IL4 signature remained a dominant feature on day 6 (Figure 4C, Supplementary Table 2D-E). A Th2 skew and IL-4 have both been associated with lung damage in COVID-19 patients (53). On day 6 there was an up-regulation of USRs primarily associated with tissue repair (Figure 4D, Supplementary Table 2F); MAPK1 (also known as ERK) and MAPK9 (54), BMP7 (55), TCF7L2 (56), and KLF4 (57, 58). B and T cell-associated USRs were also seen (Figure 4D, green circles) (59–61), consistent with development of an adaptive immune response. A relatively minor PARP1 signature was identified (Figure 4D), with PARP inhibitors being considered for COVID-19 therapy (62, 63). A strong signature associated with estrogen receptor 1 (ESR1) was observed (Figure 4D), with estrogen having anti-inflammatory properties and believed to be associated with reduced COVID-19 severity in women (64–66), and reduced SARS-CoV disease severity in female mice (46, 67). Follicle stimulating hormone (FSH) (Figure 4D) stimulates estrogen production, and inhibin subunit alpha (INHA) is involved in the negative feedback of FSH production (68). IPA Diseases and Functions analysis, Cytoscape, and GO Biological Processes analyses further supported the contention that on day 6 inflammation had abated and tissue repair was underway (Supplementary Table 2G-L).

The >31,000 gene sets available in MSigDB were used to interrogate the complete gene list (pre-ranked by fold change) for day 6 versus day 2 (Supplementary Table 2B) by GSEA. A highly significant negative enrichment for inflammatory response, and a highly significant positive enrichment for cilium development was seen (Figure 4E). Cilia in mammals are found in the lining of the respiratory epithelium. These GSEAs thus provide further support for the conclusion above, together arguing that in the hACE2-lentivirus model, day 6 post infection is characterized by inflammation resolution and tissue repair.

### Type I and III IFN signaling is required for SARS-CoV-2-induced lung inflammation

To determine what responses to SARS-CoV-2 infection are dependent on type I IFN signaling, the lentivirus system was used to analyze IFNAR^-/-^ mice lung using RNA-Seq. IFNAR^-/-^ and C57BL/6J mice were transduced with hACE2-lentivirus and were then infected with SARS-CoV-2 and lungs harvested for RNA-Seq on days 2 and 6 post-infection. The largest impact of type I IFN signaling deficiency was seen on day 2 post infection (Figure 5A), consistent with disease resolution on day 6 (Figure 4C-E). The total number of DEGs for IFNAR^-/-^ versus C57BL/6J mouse lungs on day 2 was 192, with most of these down-regulated (Figure 5A, Supplementary Table 3A-C). Viral reads were not significantly different between these two mouse strains (Figure 5A), confirming that knocking out type I IFN signaling is insufficient to significantly affect SARS-CoV-2 replication (69).

**Figure 5.**
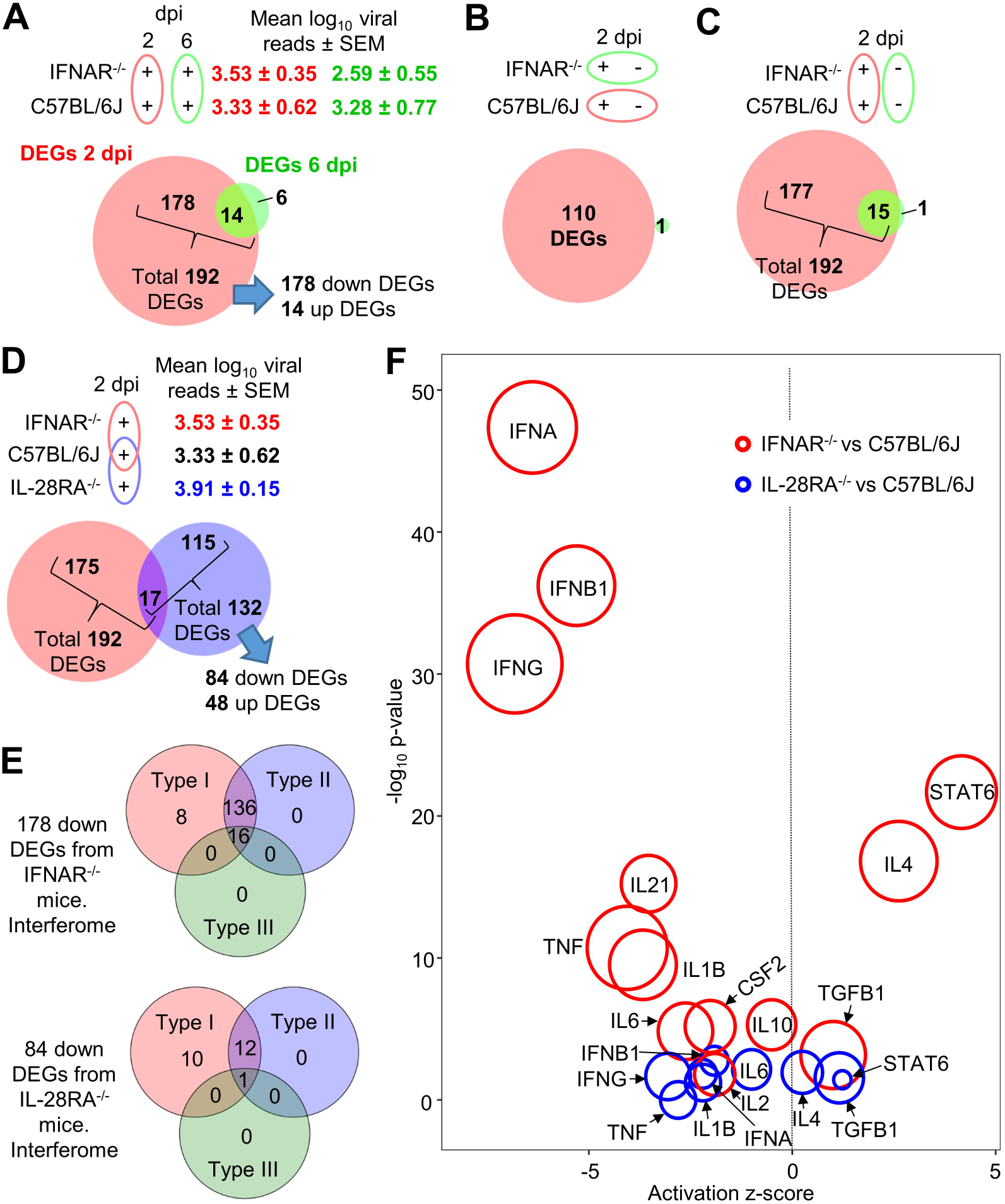
Type I and III IFN signaling is required for SARS-CoV-2-induced lung inflammation. **A)** Venn diagram for number of DEGs (FDR<0.05) between IFNAR^-/-^ versus C57BL/6J mice at day 2 or day 6 (green). Mean log_10_ SARS-CoV-2 read count in RNA-seq data ± SEM is shown (red text = day 2, green text = day 6). **B)** Venn diagram for number of DEGs (FDR<0.05) between plus versus minus hACE2-lentivirus transduction comparing C57BL/6J (red) and IFNAR^-/-^ (green) mice at day 2. **C)** Venn diagram for IFNAR^-/-^ versus C57BL/6J mice at day 2 comparing hACE2-lentivirus transduced (red) or untransduced (green) mice. **D)** Venn diagram comparing IFNAR^-/-^ versus C57BL/6J (red) and IL-28RA^-/-^ versus C57BL/6J (blue) (see Supplementary Table 3B-C and 4B-C for full gene and DEG lists) mice at day 2 (all mice had received hACE2-lentivirus transduction). Mean log_10_ SARS-CoV-2 read count in RNA-seq data ± SEM is shown (red text = IFNAR^-/-^, black text = C57BL/6J, blue text = IL-28RA^-/-^). **E)** Interferome analysis of down DEGs from ‘D’. See supplementary Table 3D and 4D for full Interferome ISG list. **F)** Signatures identified using IPA USR analysis of DEGs identified in ‘D’ for IFNAR^-/-^ versus C57BL/6J (red) and IL-28RA^-/-^ versus C57BL/6J (blue). Circle area reflects the number of DEGs associated with each USR annotation. Signatures were selected focusing on data previously identified in ‘Figure 3C’ (entire list is in Supplementary Table 3E and 4E).

We showed in Figure 3 that RNA-Seq analysis revealed 110 DEGs associated with inflammatory responses when hACE2-transduced versus untransduced C57BL/6J mouse lungs on day 2 post infection were compared. When the same comparison was made for hACE2-lentivirus transduced versus untransduced lungs in SARS-CoV-2 infected IFNAR^-/-^ mice, only 1 DEG (Kcnk5) was identified (Figure 5B). This clearly illustrated that the inflammatory responses on day 2 in C57BL/6J mice (Figure 3C) required intact type I IFN signaling.

As a control, the number of DEGs for IFNAR^-/-^ mice versus C57BL/6J in untransduced lungs was determined. Although virus replication (hACE2-lentivirus transduced lung) provided 192 DEGs, virus inoculation (untransduced lung) provided only 16 DEGs (Figure 5C). Thus again, virus inoculation had a limited effect on these analyses, with DEGs largely associated with virus infection.

Type III IFN is thought to act as a first-line of defense against infection at epithelial barriers, with the more potent and inflammatory type I IFN response kept in reserved in the event the first line of defense is breached (70, 71). As type III IFN signaling has been implicated in reducing SARS-CoV-2 infection *in vitro* (72), the aforementioned analyses on day 2 was repeated in IL-28RA^-/-^ mice. Viral reads were not significantly different between these two mouse strains (Figure 5D), confirming that knocking out type III IFN signaling was insufficient to significantly affect SARS-CoV-2 replication (69, 73). The DEG list for IL-28RA^-/-^ versus C57BL/6J mice comprised 132 genes (Supplementary Table 4A-C), of which 84 were down-regulated (Figure 5D). Perhaps surprisingly, the DEGs were mostly different from that seen in IFNAR^-/-^ mice (Figure 5D), arguing that the ISGs stimulated by type I and III IFNs are largely distinct.

The majority (90%) of the 178 down-regulated DEGs in IFNAR^-/-^ versus C57BL/6J mice on day 2 post infection were ISGs as defined by Interferome (using *in vivo* and mouse only settings) (Figure 5E, top and Supplementary Table 3D). Of the 84 down-regulated genes in IL-28RA^-/-^ lungs, only 27% were identified as ISGs (Figure 5E, bottom and Supplementary Table 4D). However, there are no annotations in the Interferome database for type III ISGs in mice, and changing the settings to include human data (which is also under-annotated for type III IFN) did not increase the number of type III ISGs in our search using down DEGs from IL-28RA^-/-^ lungs.

To determine the pathways regulated by type I and III IFN in response to SARS-CoV-2 infection, IPA USR analyses was performed on DEG lists from day 2 for hACE2-lentivirus transduced lungs for IFNAR^-/-^ versus C57BL/6J (192 DEGs) and IL-28RA^-/-^ versus C57BL/6J (132 DEGs). IPA USR signatures associated with pro-inflammatory cytokine signaling were significantly reduced in both IFNAR^-/-^ and, to a lesser extent, IL-28RA^-/-^ mice when compared to C57BL/6J mice (Figure 5F, full lists in Supplementary Table 3E and 4E). The USR annotations for type I and III IFN receptor deficiencies were largely similar (Figure 5F), although they arose from a largely distinct set of DEGs (Figure 5D). This likely reflects expression of type I and type III IFN by different cell types with differing response profiles to the same cytokines (70).

Taken together these results illustrate that while there was no significant effects on viral replication, lungs from type I and type III IFN receptor knockout mice both had a significantly blunted acute pro-inflammatory response following SARS-CoV-2 infection. The importance of type I IFN for inflammation was reported previously using the AAV-hACE2 system (18).

### Immunization with infectious or UV-inactivated SARS-CoV-2 protects against virus infection

To evaluate the utility of the hACE2-lentivirus mouse model of SARS-CoV-2 replication for vaccine studies, C57BL/6J mice were immunized twice subcutaneously (s.c.) with either infectious SARS-CoV-2 or UV-inactivated SARS-CoV-2 (74) formulated with adjuvant (Figure 6A). Serum SARS-CoV-2 neutralization titers were determined post-boost, and revealed all mice had significant serum neutralization titers (Figure 6B), comparable with the higher end levels seen in people with past SARS-CoV-2 infections (75). On day 2 post challenge all mice were sacrificed. All mouse lungs had detectable hACE2 mRNA by RT-qPCR, with levels higher in unvaccinated mice (Figure 6C), likely due to immune cell infiltrates (see below) diluting hACE2 mRNA. Mice immunized with either infectious or UV-inactivated SARS-CoV-2 showed significant reductions in viral RNA levels by RT-qPCR (Figure 6D) in tissue titrations (Figure 6E) and by RNA-Seq read counts (Figure 6F). These immunizations thus protected mice against significant viral infections and demonstrated the utility of hACE2-lentivirus transduced mice as a model for vaccine evaluation.

**Figure 6.**
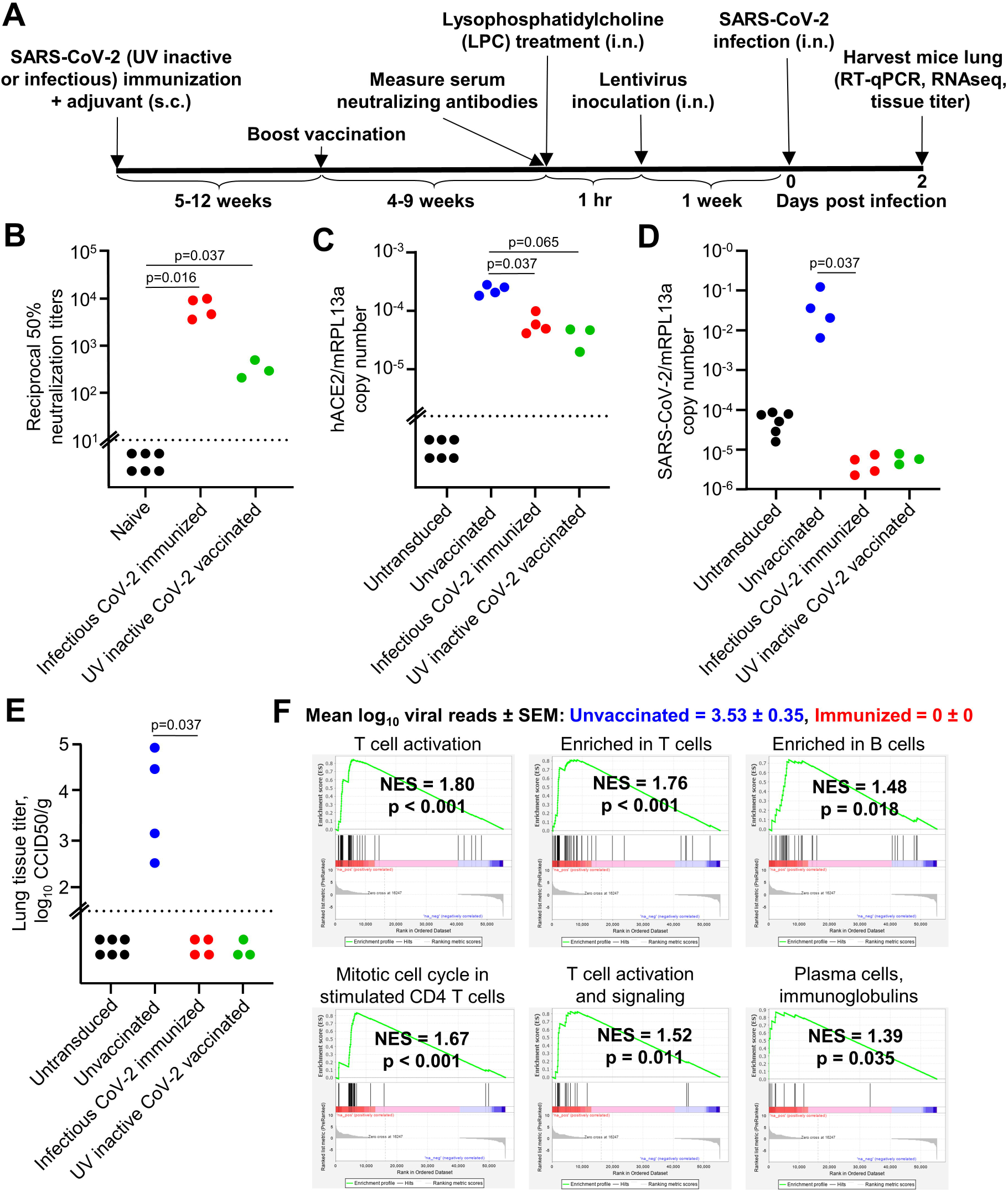
Immunization with infectious or UV-inactivated SARS-CoV-2 protects against virus infection. **A)** Timeline of mice immunization, lentivirus transduction and SARS-CoV-2 infection. **B)** Reciprocal 50% neutralization titers of naïve mice and infectious or UV SARS-CoV-2 immunized mice. Horizontal dotted line represents the limit of detection (1 in 10). Statistics are by Kolmogorov Smirnov test. **C)** RT-qPCR of mice lung RNA using primers for hACE2 introduced by lentivirus transduction normalized to mRPL13a levels. Data is individual mice and is expressed as RNA copy number calculated against a standard curve for each gene. Horizontal line indicates cut-off for reliable detection, with all hACE2-negative mice falling below this line. Statistics are by Kolmogorov Smirnov test. **D)** RT-qPCR of mice lung RNA using primers for SARS-CoV-2 E gene normalized to mRPL13a levels. Data is individual mice and is expressed as RNA copy number calculated against a standard curve for each gene. Statistics are by Kolmogorov Smirnov test. Unvaccinated versus UV-inactive CoV-2 vaccinated reaches significance by Kruskal-Wallis test (p=0.034). **E)** Titer of SARS-CoV-2 in mice lung determined using CCID_50_ assay of lung homogenate. Horizontal line indicates the limit of detection of 1.57 log_10_ CCID_50_/g. Statistics are by Kolmogorov Smirnov test. Unvaccinated versus UV-inactive CoV-2 vaccinated reaches significance by Kruskal-Wallis test (p=0.028). **F)** Mean log_10_ SARS-CoV-2 read count at day 2 post-infection in RNA-seq data ± SEM is shown (blue text = unvaccinated mice, red text = infectious SARS-CoV-2 immunized mice). GSEA for Blood Transcription Modules (BTMs) from Li et al. (76) against the pre-ranked all gene list comparing unvaccinated versus infectious SARS-CoV-2 immunized (s.c) mice lung at day 2 (all mice had hACE2-lentivirus transduction). Selected BTM modules with a positive NES score and p < 0.05 are shown (full list in Supplementary Table 5). See Supplementary Table 5 for full gene and DEG lists and full downstream analyses lists.

Lungs harvested on day 2 were also analyzed by RNA-Seq (Supplementary Table 5A-B), with 1376 DEGs identified for infectious virus immunization versus unimmunized (DEGs in Supplementary Table 5C). The DEGs were analyzed by IPA canonical pathways, USR analysis, and the Diseases and Functions feature, which all showed infiltration signatures, dominated by T cell annotations (Supplementary Table 5D-F). This was confirmed by GSEA of human blood transcription modules (76) against the pre-ranked (by fold change) gene list for immunized versus unvaccinated mice (Supplementary Table 5G). This showed the expected enrichment of T cell and B annotations in immunized mice (Figure 6F), and illustrated that protection against virus replication in lungs of immunized mice was associated with a significant infiltration of lymphocytes. This aligns with studies that suggest COVID-19 patients that induce an early SARS-CoV-2-specific T cell response have rapid viral clearance and mild disease (77).

## DISCUSSION

Herein we describe a hACE2-lentivirus transduction system that allows the use of C57BL/6J and GM mice for SARS-CoV-2/COVID-19 research. Virus titer in hACE2-lentivirus transduced mice lung were comparable to other studies using mice transduced with Ad5 or AAV expressing hACE2 (10, 17, 18). These transduction methodologies avoid the fulminant brain infection seen in K18-hACE2 mice, with infection of this organ the major contributor to mortality in the K18-hACE2 model (78, 79). The key advantage of lentiviral over adenoviral systems is the integration of the lentiviral payload into the host cell genome, thereby providing long-term stable expression. The hACE2-lentivirus thus provides a new option for studying SARS-CoV-2 infection that can be readily applied using standard lentivirus technology. We characterize the lentivirus-hACE2 model of SARS-CoV-2 infection in C57BL/6J mice, and illustrate the use of lentivirus-hACE2 to assess the role of type I and type III IFNs in virus control and inflammatory responses using IFNAR^-/-^ and IL-28RA^-/-^ mice, respectively. We also illustrate the use of C57BL/6J mice transduced with lentivirus-hACE2 for SARS-CoV-2 vaccine evaluation.

The lentivirus-hACE2 C57BL/6J mouse model of SARS-CoV-2 infection shares many cytokine signatures with those seen in human COVID-19 patients; IL-2, IL-10, IL-6, TNFα, IL-4, IL-1β, IFNγ and CSF3 signatures were found in SARS-CoV2-infected lentivirus-hACE2 transduced C57BL/6J mice on day 2 post infection, and elevated levels of these cytokines are found in COVID-19 patients (80, 81). Several of these cytokines have been implicated as mediators of COVID-19; for instance, IL-6 (82–84), IL-4 (53), IL-10 (85) and IFNγ (86). On day 6 post infection the lentivirus-hACE2 C57BL/6J mouse model showed a series of signatures associated with tissue repair. Thus our RNA-Seq analyses illustrated that this model involves an acute inflammatory disease followed by resolution and tissue repair.

hACE2-lentivirus transduction in IFNAR^-/-^ and IL-28RA^-/-^ mice was used to confirm that type I IFN signaling (18) and show that type III IFN signaling are involved in driving inflammatory responses. Virus replication was not significantly affected in IFNAR^-/-^ and IL-28RA^-/-^ mice, largely consistent with other studies using different model systems, which indicate that knocking out both pathways (STAT1^-/-^, STAT2^-/-^, or combined IFNAR^-/-^/IL-28RA^-/-^) is required before an impact on virus replication is seen (18, 69, 73, 87). A high level of cross-compensation is thus implicated, as is described in influenza infection (71). *In vitro* experiments show that SARS-CoV-2 replication is sensitive to type I and III IFNs, although infection does not stimulate particularly high levels of IFN (72, 88, 89). A number of SARS-CoV-2 encoded proteins inhibit IFN production and signaling (83, 90, 91). In infected cells, such inhibition of signaling would limit the antiviral activity of IFNs, as well as limiting the positive feedback amplification that is needed for high level IFN production (92). IFN therapies have been proposed for COVID-19 patients (93) and such treatment may induce an antiviral state in uninfected cells, thereby perhaps limiting viral spread if given prophylactically or early during the course of infection (94). However, caution might be warranted for such therapy during later stage COVID-19 as type I and III IFN signaling are shown herein to drive inflammatory responses during SARS-CoV-2 infection, and are also known to have pro-inflammatory activities in other settings (92).

Herein we also illustrate the use of the hACE2-lentivirus system to study the role of ACE2 residues on SARS-CoV-2 replication *in vitro*. Only four “mouse-to-human” changes in mACE2 (N30D, F83Y, N31K, H353K) enabled full SARS-CoV-2 replication to levels similar to those seen for infection of hACE2-expressing cells. These results suggest a CRISPR mouse with these substitutions would be able to support SARS-CoV-2 infection and provide a model of COVID-19. Remarkably, a minimum of only two substitutions in mACE2 (N31K, H353K) were sufficient for partial restoration of SARS-CoV-2 infection. Changing the same resides in hACE2 to their mouse equivalents (K31N, K353H) did not affect virus replication compared to WT hACE2. Thus N31K and H353K allows binding of SARS-CoV2 to mACE2, but K31N and K353H does not impair binding to hACE2. Both K31N and K353H substitutions were shown to be deleterious for SARS-CoV2 binding by *in silico* analyses. However, the same analyses implicated >25 other residues in hACE2 binding (31), with many of these potentially able to compensate for the loss of native residues at K31 and K353.

In conclusion, we describe the development of a mouse model using intranasal hACE2-lentivirus transduction to sensitize mice lung to SARS-CoV-2 infection, which recapitulated cytokine signatures seen in other mouse models and COVID-19. We demonstrate broad applicability of this new mouse model in vaccine evaluation, GM mice infection, and *in vitro* evaluation of ACE2 mutants.

## ACKNOWLEDGEMENTS

We thank Dr I Anraku for his assistance in managing the PC3 (BSL3) facility at QIMR Berghofer MRI. We thank Dr Alyssa Pyke and Mr Fredrick Moore (Queensland Health, Brisbane) for providing the SARS-CoV-2 isolate. We thank Dr. David Harrich for providing the lentivirus vector system. We thank Dr. Simon Phipps for providing the IL-28RA^-/-^ mice. We thank Monash Genome Modification Platform for providing the plasmid containing the mouse-codon optimized human ACE2 gene. We thank Clive Berghofer and Lyn Brazil (and many others) for their generous philanthropic donations to support SARS-CoV-2 research at QIMR Berghofer MRI. A.S. holds an Investigator grant from the National Health and Medical Research Council (NHMRC) of Australia (APP1173880).

## AUTHOR CONTRIBUTIONS

Conceptualization, D.J.R.; Methodology, D.J.R. and A.S.; Formal analysis, D.J.R., A.S., and T.D.; Investigation, D.J.R., T.T.L., K.Y., B.T.; Data curation, D.J.R., A.S., and T.D.; Writing – original draft, D.J.R.; Writing – review and editing, A.S. and D.J.R.; Visualization, D.J.R., A.S., T.D., and C.B; Supervision, D.J.R. and A.S., Project administration, D.J.R. and A.S.; Funding acquisition, A.S. and D.J.R.

## DATA AVAILABILITY

All raw sequencing data (fastq files) are available from the Sequence Read Archive (SRA), BioProject accession: PRJNA701678. All other data is available within the paper and supporting information files.

## MATERIALS and METHODS

### Ethics statement

All mouse work was conducted in accordance with the “Australian code for the care and use of animals for scientific purposes” as defined by the National Health and Medical Research Council of Australia. Mouse work was approved by the QIMR Berghofer Medical Research Institute animal ethics committee (P3600, A2003-607). For intranasal inoculations, mice were anesthetized using isoflurane. Mice were euthanized using CO_2_ or cervical dislocation.

### Cell lines and SARS-CoV-2 culture

Vero E6 (C1008, ECACC, Wiltshire, England; Sigma Aldridge, St. Louis, MO, USA), HEK293T (a gift from Michel Rist, QIMR Berghofer), Lenti-X 293T (Takara Bio), AE17 (a gift from Delia Nelson, Faculty of Health Sciences, Curtin Medical School), and NIH-3T3 (American Type Culture Collection, ATCC, CRL-1658) cells were cultured in medium comprising DMEM for Lenti-X 293T cells or RPMI1640 for all others (Gibco) supplemented with 10% fetal calf serum (FCS), penicillin (100◻IU/ml)/streptomycin (100◻μg/ml) (Gibco/Life Technologies) and L-glutamine (2 mM) (Life Technologies). Cells were cultured at 37°C and 5% CO_2_. Cells were routinely checked for mycoplasma (MycoAlert Mycoplasma Detection Kit MycoAlert, Lonza) and FCS was assayed for endotoxin contamination before purchase (96). The SARS-CoV-2 isolate was kindly provided by Queensland Health Forensic & Scientific Services, Queensland Department of Health, Brisbane, Australia. The virus (hCoV-19/Australia/QLD02/2020) was isolated from a patient and sequence deposited at GISAID (https://www.gisaid.org/; after registration and login, sequence can be downloaded from https://www.epicov.org/epi3/frontend#1707af). Virus stock was generated by infection of Vero E6 cells at multiplicity of infection (MOI)≈0.01, with supernatant collected after 2-3 days, cell debris removed by centrifugation at 3000 x g for 15 min at 4°C, and virus aliquoted and stored at −80°C. Virus titers were determined using standard CCID_50_ assays (see below). The virus was determined to be mycoplasma free using co-culture with a non-permissive cell line (i.e. HeLa) and Hoechst staining as described (97).

### CCID_50_ assays

Vero E6 cells were plated into 96 well flat bottom plates at 2×10^4^ cells per well in 100 µl of medium. For tissue titer, tissue was homogenized in tubes each containing 4 ceramic beads twice at 6000 x g for 15 seconds, followed by centrifugation twice at 21000 x g for 5 min before 5 fold serial dilutions in 100 µl RPMI1640 supplemented with 2% FCS. For cell culture supernatant, 10 fold serial dilutions were performed in 100 µl RPMI1640 supplemented with 2% FCS. 100 µl of serially diluted samples were added to Vero E6 cells and the plates cultured for 5 days at 37°C and 5% CO_2_. The virus titer was determined by the method of Spearman and Karber (a convenient Excel CCID_50_ calculator is available at https://www.klinikum.uni-heidelberg.de/zentrum-fuer-infektiologie/molecular-virology/welcome/downloads).

### Lentivirus cloning

ACE2 genes were cloned into pCDH-EF1α-MCS-BGH-PGK-GFP-T2A-Puro Cloning and Expression Lentivector (System Biosciences, catalogue number CD550A-1), where the EF1α promoter was replaced with the full length EF1α using NheI and ClaI restriction enzymes (New England Biolabs). Human ACE2 coding sequence (codon optimized for mouse) was cloned into the pCDH lentivector with PCR fragments amplified using Q5® High-Fidelity 2X Master Mix (New England Biolabs) and the following primers; vector backbone (pCDH amplified with Forward 5’-TAAATCGGATCCGCGG -3’ and Reverse 5’-AATTCGAATTCGCTAGC) and hACE2 insert (Forward 5’-GCTAGCGAATTCGAATTATGAGCAGCAGCTCTTGGC -3’ and Reverse 5’-CCGCGGATCGCATTTATCAGAAGCTTGTCTGCACGT -3’). The pCDH fragment was digested with DpnI (New England Biolabs) and was purified using QIAquick Gel Extraction Kit (QIAGEN), as was the hACE2 fragment. Fragments were recombined using NEBuilder HiFi DNA Assembly Cloning Kit as per manufacturer instructions. This was transformed into NEB® 10-beta Competent E. coli (High Efficiency) (New England Biolabs) as per manufacturer instructions and plated on ampicillin agar plates overnight. Colony PCR was performed using Q5 High-Fidelity 2X Master Mix (New England Biolabs) and the hACE2 insert primers to identify a positive colony, which was grown in LB broth with ampicillin and plasmid was purified using NucleoBond Xtra Midi kit (Machery Nagel).

All ACE2 mutant coding sequences were ordered from Twist Bioscience/Decode Science containing EcoRI upstream and BamHI downstream. Fragments were amplified using the using Q5® High-Fidelity 2X Master Mix (New England Biolabs) and the following primers; Forward 5’-CCTGACCTTAGCGAATTCATG -3’ and Reverse 5’-ACCTAGCCTCGCGGATC -3’. PCR fragments were purified using Monarch DNA Gel Extraction Kit (New England Biolabs), and were digested with EcoRI and BamHI (New England Biolabs), as was the pCDH vector before purification again with Monarch® DNA Gel Extraction Kit (New England Biolabs). The ACE2 fragment was ligated with pCDH vector using T4 DNA Ligase (New England Biolabs) as per manufacturer instructions and then transformed into NEB® 10-beta Competent E. coli (High Efficiency) (New England Biolabs) as per manufacturer instructions and plated on ampicillin agar plates overnight. Colonies were grown in LB broth with ampicillin and plasmid was purified using NucleoBond Xtra Midi kit (Machery Nagel). Plasmids were confirmed by PCR with the cloning primers above and fragments were gel purified and confirmed by Sanger sequencing with the forward primer.

### Lentivirus production, titration and cell line transduction

ACE2 lentivirus was produced by co-transfection of HEK293T or Lenti-X HEK293T cells with the pCDH-ACE2 plasmid, VSV-G and Gag-Pol using Lipofectamine 2000 Reagent (Thermo Fisher Scientific) or Xfect Transfection Reagent (Takara Bio) as per manufacturer instructions. Supernatant was collected 2 days after transfection and centrifuged at 500 x g for 10 min, and this was concentrated using Amicon Ultra-15 Centrifugal Filter Units with 100 kDa cutoff (Merck Millipore) as per manufacturer instructions. Lentivirus was titrated by serial dilution of lentivirus and incubating with 5000 HEK293T cells in 96 well plates with 8 µg/ml polybrene for 3 days followed by GFP detection by flow cytometry (BD Biosciences LSRFortessa). Transduction units (TU) per ml was calculated by percentage of 5000 cells that are GFP positive multiplied by the dilution. For example if 2 µl lentivirus gives 5% GFP positive cells, the TU/ml is ((5/100)*5000) * (1000/2) = 125,000 TU/ml. The p24 equivalent (ng/ml) was measured using Lenti-X GoStix Plus (Takara Bio) as per manufacturer instructions.

Cell lines were transduced by incubating with lentivirus and 8 µg/ml polybrene for 2-3 days followed by 5 µg/ml puromycin treatment until most resistant cells expressed GFP. Cells were then infected with SARS-CoV-2 at MOI 0.1 for 1 hr at 37°C, cells were washed with PBS and media replaced. Culture supernatant was harvested at the indicated timepoints and titered by CCID_50_ assay as described.

### SARS-CoV-2 spike RBD:ACE2 mutagenesis modeling

PyMOL v4.60 (Schrodinger) was used for mutagenesis of the crystal structure of SARS-CoV-2 spike receptor-binding domain bound with ACE2 from the protein data bank (6M0J) (95). The side chain orientation (rotamer) with highest frequency of occurrence in proteins was chosen for all mutations. Polar interactions (blue lines) or any interactions within 3.5Å (yellow lines) were shown for selected residues. The K417N and N501Y mutations in the UK variant (B.1.1.7) and RSA variant (B.1.351) were introduced (38).

### UV-inactivated SARS-CoV-2 vaccine preparation and mice immunization

120 ml of SARS-CoV-2 was inactivated with at least 7650 J/m^2^ UVC in a UVC 500 Ultraviolet Crosslinker (Hoefer). Virus was determined to be inactivated by incubation with Vero E6 cells for 4 days without development of CPE. UV-inactive SARS-CoV-2 was then partially-purified using a 20% sucrose cushion centrifuged at 175,000 x g for 3-4 hr using SW32Ti rotor (Beckman Coulter). The sample that has passed through the sucrose was collected and concentrated using Amicon Ultra-15 Centrifugal Filter Units with 100 kDa cutoff (Merck Millipore). For infectious SARS-CoV-2 immunization, virus was also concentrated using Amicon Ultra-15 Centrifugal Filter Units with 100 kDa cutoff (Merck Millipore). Mice were injected s.c. in the base of the tail with 100 µl of UV-inactive or infectious SARS-CoV-2 with 25 µg adjuvant in 50 µl made with the same ingredients as AS01 (98) as described previously (99). A boost was given 5-6 weeks after prime, and mice were bled to measure serum neutralizing titers 4-9 weeks after boost.

### Neutralization assay

Mouse serum was heat inactivated at 56°C for 30 min and incubated with 100 CCID_50_ SARS-CoV-2 for 2 hr at 37°C before adding 10^5^ Vero cells/well in a 96 well plate to 200 µl. After 4 days cells were fixed and stained by adding 50 µl formaldehyde (15% w/v) and crystal violet (0.1% w/v) (Sigma-Aldrich) overnight. The plates were washed in tap water, dried overnight and 100 µl/well of 100% methanol added to dissolve the crystal violet and the OD was read at 595 nm using a 96-well plate reader (Biotek Synergy H4). The 50% neutralizing titers were interpolated from optical density (OD) versus dilution plots.

### Mice intranasal lentivirus transduction and SARS-CoV-2 infection

C57BL/6J, IFNAR^-/-^ (100) (originally provided by P. Hertzog, Monash University, Melbourne, VIC, Australia) and IL-28RA^-/-^ mice (71, 101) were kindly provided by Bristol-Myers Squibb (102) and bred in-house at QIMRB. Female mice were 8 weeks to 1 year old (age matched between groups) at the start of the experiment. Mice were anesthetized using isoflurane and 4 µl of 1% L-α-Lysophosphatidylcholine from egg yolk (Sigma Aldrich) in water was administered intranasally. After 1 hour mice were inoculated intranasally with approximately 2×10^4^ TU of hACE2-pCDH lentivirus in 50 µl, and 1 week later challenged with 10^5^ CCID_50_ SARS-CoV-2 intranasally in 50 µl. Mice were sacrificed by cervical dislocation at day 2, 4 or 6 and lungs were collected. Right lung was immediately homogenized in tubes each containing 4 beads twice at 6000 x g for 15 seconds, and used in tissue titration as described above. Left lung was placed in RNAprotect Tissue Reagent (QIAGEN) at 4°C overnight then −80°C.

K18-hACE2 mice (6) were purchased from The Jackson Laboratory and bred in-house at QIMRB with C57BL/6J mice. Mice were genotyped using Extract-N-Amp Tissue PCR Kit (Sigma Aldrich) according to manufacturer instructions with the following primers; Forward 5’-CTTGGTGATATGTGGGGTAGA-3’ and Reverse 5’-CGCTTCATCTCCCACCACTT-3’. hACE2 positive mice were infected with 5×10^4^ SARS-CoV-2 i.n. as above and lung RNA was harvested at day 4 for RNA-seq.

### RT-qPCR

Mice lung was transferred from RNAlater to TRIzol (Life Technologies) and was homogenized twice at 6000 x g for 15 sec. Homogenates were centrifuged at 14,000 × g for 10 min and RNA was isolated as per manufacturer’s instructions. cDNA was synthesized using ProtoScript II First Strand cDNA Synthesis Kit (New England Biolabs) and qPCR performed using iTaq Universal SYBR Green Supermix (Bio-Rad) as per manufacturer instructions with the following primers; SARS-CoV-2 E Forward 5’-ACAGGTACGTTAATAGTTAATAGCGT -3’ and Reverse 5’- ATATTGCAGCAGTACGCACACA, hACE2 Forward 5’- GATCACGATTCCCAGGACG -3’ and Reverse 5’- TCCGGCTGAACGACAACTC -3’, mRPL13a (103) Forward 5’- GAGGTCGGGTGGAAGTACCA -3’ and Reverse 5’- TGCATCTTGGCCTTTTCCTT -3’. PCR fragments of SARS-CoV-2 E, ACE2 and mRPL13a using the same primers as above were gel purified and 10-fold serial dilutions of estimated copy numbers were used as standards in qPCR to calculate copies in samples reactions. SARS-CoV-2 E and ACE2 copies were normalized by mRPL13a copy number in each reaction. qPCR reactions were performed in duplicate and averaged to determine the copy number in each sample.

### RNA-seq

TRIzol extracted lung RNA was treated with DNase (RNase-Free DNAse Set (Qiagen)) followed by purification using RNeasy MinElute Cleanup Kit (QIAGEN) as per manufacturer instructions. RNA concentration and quality was measured using TapeStation D1K TapeScreen assay (Agilent). cDNA libraries were prepared using the Illumina TruSeq Stranded mRNA library prep kit and the sequencing performed on the Illumina Nextseq 550 platform generating 75bp paired end reads. Per base sequence quality for >90% bases was above Q30 for all samples. The quality of raw sequencing reads was assessed using FastQC (104) (v0.11.8) and trimmed using Cutadapt (105) (v2.3) to remove adapter sequences and low-quality bases. Trimmed reads were aligned using STAR (106) (v2.7.1a) to a combined reference that included the mouse GRCm38 primary assembly and the GENCODE M23 gene model (107), SARS-CoV-2 isolate Wuhan-Hu-1 (NC_045512.2; 29903 bp) and the human ACE2 mouse codon optimized sequence (2418 bp). Mouse gene expression was estimated using RSEM (108) (v1.3.0). Reads aligned to SARS-CoV-2 and hACE2 were counted using SAMtools (109) (v1.9). Differential gene expression in the mouse was analyzed using EdgeR (3.22.3) and modelled using the likelihood ratio test, glmLRT().

### Analyses of K18-hACE2 and Ad5-hACE2 RNA-seq data

RNA-seq datasets generated from the Winkler et al. study (9), and Sun et al. study (43) were obtained from the Gene Expression Omnibus (GSE154104 and GSE150847 respectively) and trimmed using Cutadapt (v2.3). Trimmed reads were aligned using STAR (v2.7.1a) to a combined reference that included the mouse GRCm38 primary assembly and the GENCODE M23 gene model, SARS-CoV-2 isolate Wuhan-Hu-1 (NC_045512.2; 29903 bp) and the human ACE2 transcript variant 1 (NM_001371415.1; 3339 bp). Mouse gene expression was estimated using RSEM (v1.3.0). Reads aligned to SARS-CoV-2 and hACE2 were counted using SAMtools (v1.9). Differential gene expression in the mouse was analyzed using EdgeR (3.22.3) and modelled using the likelihood ratio test, glmLRT().

### Pathway Analysis

Up-Stream Regulators (USR), Diseases and Functions and canonical pathways enriched in differentially expressed genes in direct and indirect interactions were investigated using Ingenuity Pathway Analysis (IPA) (QIAGEN).

### Network Analysis

Protein interaction networks of differentially expressed gene lists were visualized in Cytoscape (v3.7.2) (110). Enrichment for biological processes, molecular functions, KEGG pathways and other gene ontology categories in DEG lists was elucidated using the STRING database (111) and GO enrichment analysis (112).

### Gene Set Enrichment Analysis

Preranked GSEA (113) was performed on a desktop application (GSEA v4.0.3) (http://www.broadinstitute.org/gsea/) using the “GSEAPreranked” module. Gene sets from the supplemental materials from the Winkler et al. (9), Blanco-Melo et al. (51) and Wu et al. (52) (filename.GMT) studies were investigated for enrichment in our pre-ranked all gene list for SARS-CoV-2 infected day 2 mice lung comparing plus versus minus ACE2 (Supplementary Table 2B). Li et al. (76) blood transcription modules (BTM_for_GSEA_20131008.gmt, n = 346) was also used in GSEA to determine enrichment in our ‘immunized’ versus unvaccinated mice lung at day 2 (Supplementary Table 5B). The complete Molecular Signatures Database (MSigDB) v7.2 gene set collection (31,120 gene sets) (msigdb.v7.2.symbols.gmt: https://www.gsea-msigdb.org/gsea/msigdb/download_file.jsp?filePath=/msigdb/release/7.2/msigdb.v7.2.symbols.gmt) was used to run GSEAs on pre-ranked gene list for plus versus minus ACE2 at day 2 (Supplementary Table 1B), and day 6 versus day 2 (Supplementary Table 2B). Gene ontology (114, 115) and Enrichr (116, 117) web-tools were also consulted.

### Interferome

The indicated DEG lists were entered into Interferome (www.interferome.org) (118) with the following parameters: *In Vivo*, Mus musculus, fold change 2 (up and down).

### Statistics

Statistical analyses of experimental data were performed using IBM SPSS Statistics for Windows, Version 19.0 (IBM Corp., Armonk, NY, USA). The t-test was used when the difference in variances was <4, skewness was >2 and kurtosis was <2. Otherwise, the non-parametric Kolmogorov–Smirnov test or Kruskal-Wallis test was used. Repeated Measures Two-Way ANOVA was used where indicated.

## SUPPPLEMENTAL INFORMATION

**Supplementary Figure 1.**
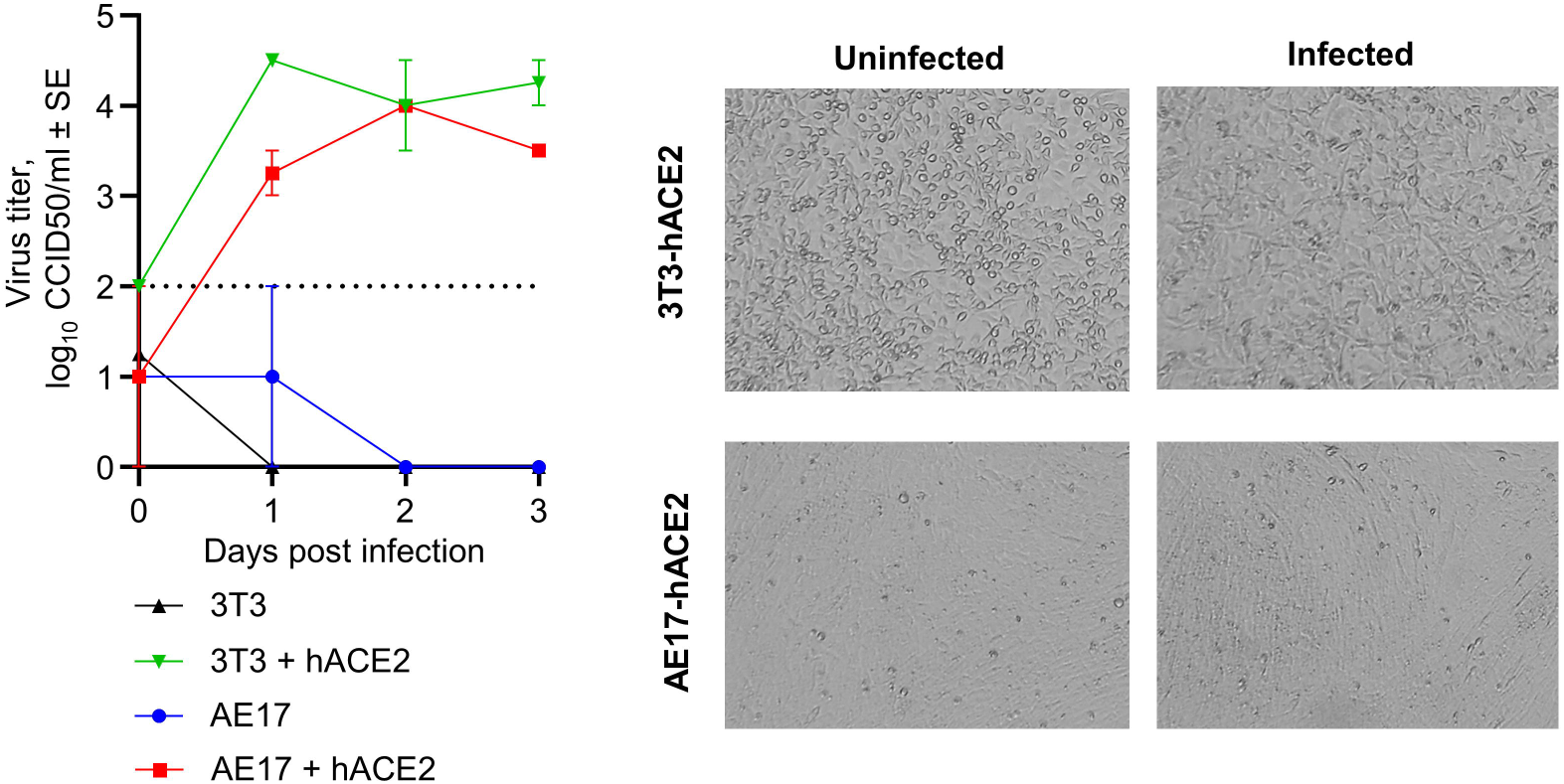
Replication of SARS-CoV-2 in hACE2-lentivirus transduced mouse cell lines. Growth kinetics of SARS-CoV-2 over a three day time course in untransduced or hACE2-lentivirus transduced 3T3 or AE17 cells infected at MOI=0.1. Data is the mean of triplicate wells and error bars represent SEM. Images of cells were taken using an inverted light microscope at day 3 post-infection and are representative of triplicate wells.

**Supplementary Figure 2.**
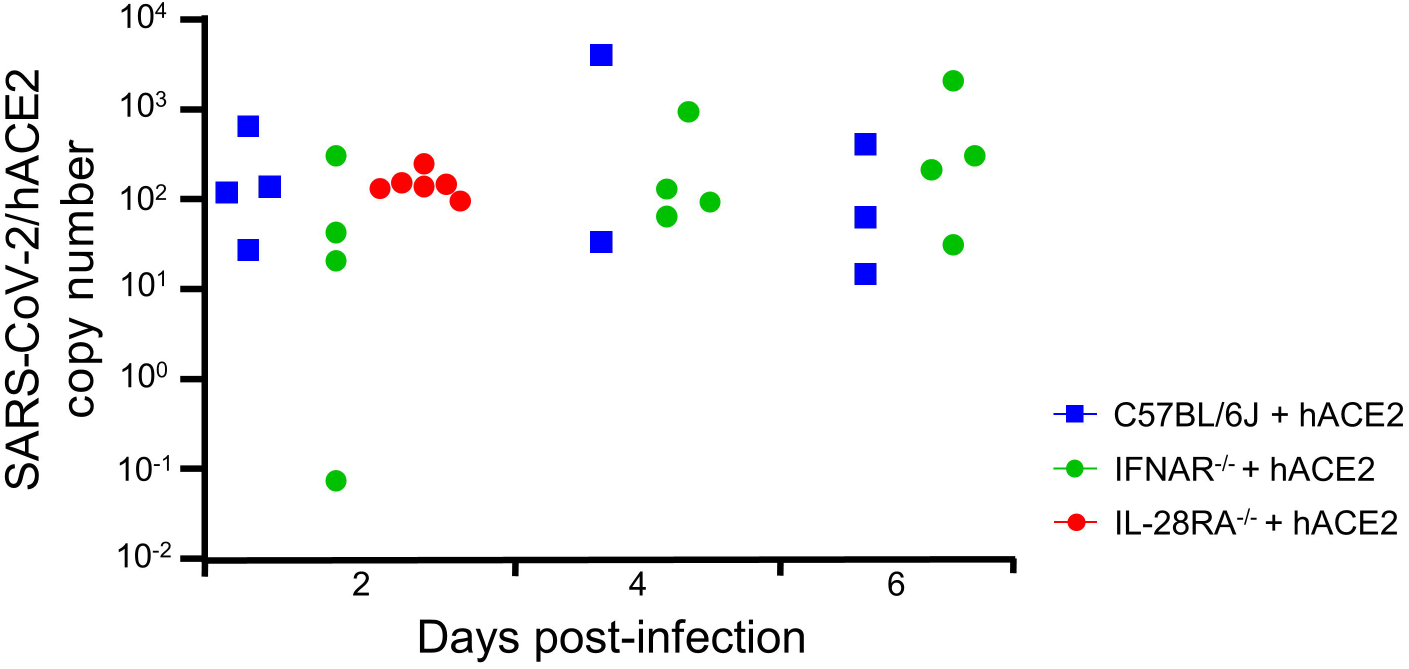
RT-qPCR of mice lung RNA using primers for SARS-CoV-2 normalized to hACE2 introduced by lentivirus transduction. Data is individual mice from Figure 2E normalised to 2C, and is expressed as RNA copy number calculated against a standard curve for each gene.

**Supplementary Figure 3.**
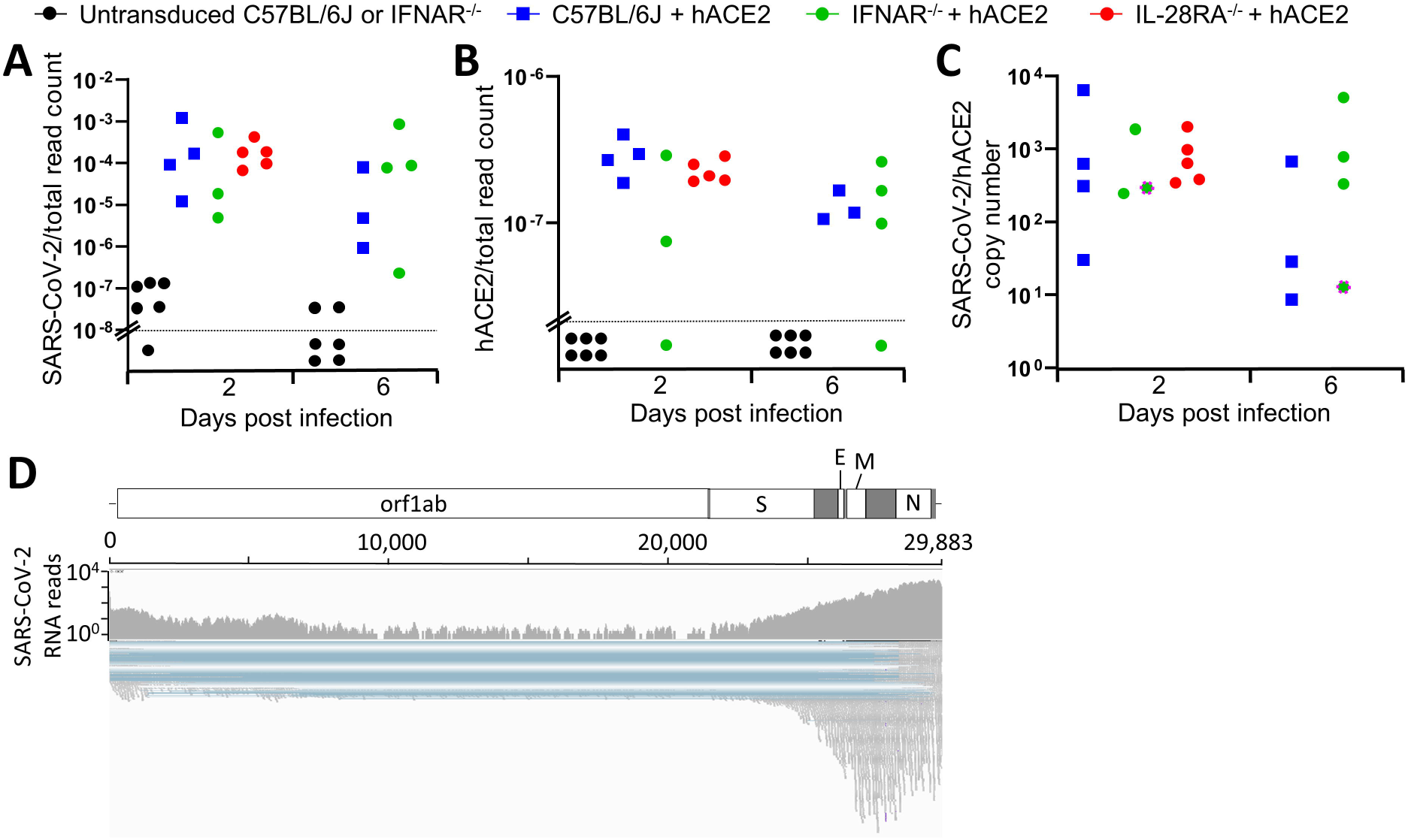
SARS-CoV-2 and hACE2 read counts in RNA-Seq data. RNA-Seq was performed on mice lung from the same mice as in ‘Figure 2’. **A)** SARS-CoV-2 read counts normalized to total read count. Data points below the horizontal dotted line had read counts of zero. **B)** hACE2 read counts normalised to total read count. Data points below the horizontal dotted line had read counts of zero. **C)** SARS-CoV-2 read count normalised to hACE2 read count. Circle with pink dotted outline had hACE2 read count of 0, so the value was set to the SARS-CoV-2 read count (not normalised). **D)** SARS-CoV-2 reads aligned to reference genome viewed in Integrative Genome Viewer (IGV). Mice with hACE2 displayed reads mapped across the entire genome, with higher counts for structural gene sub-genomic RNA as also evident in hACE2-adenoviral vector transduced cell lines (51).

**Supplementary Table 1. RNA-Seq gene lists and downstream bioinformatic analyses of responses at day 2 post-infection.**

**Supplementary Table 2. RNA-Seq gene lists and downstream bioinformatic analyses of responses at day 6 post-infection.**

**Supplementary Table 3. RNA-Seq gene lists and downstream bioinformatic analyses of responses in IFNAR^-/-^ versus C57BL/6J mice at day 2 post-infection.**

**Supplementary Table 4. RNA-Seq gene lists and downstream bioinformatic analyses of responses in IL-28RA^-/-^ versus C57BL/6J mice at day 2 post-infection.**

**Supplementary Table 5. RNA-Seq gene lists and downstream bioinformatic analyses of responses in immunized versus unvaccinated mice at day 2 post-infection.**

